# Feedback and feedforward control are differentially delayed in cerebellar ataxia

**DOI:** 10.1101/2025.02.09.637327

**Authors:** Di Cao, Michael G.T. Wilkinson, Amy J. Bastian, Noah J. Cowan

**Affiliations:** Johns Hopkins University; Johns Hopkins University, Kennedy Krieger Institute

## Abstract

Damage to the cerebellum can cause ataxia, a condition associated with impaired movement coordination. Typically, coordinated movement relies on a combination of anticipatory mechanisms (specifically, feedforward control) and corrective mechanisms (embodied by feedback control). Here, we show that in 3D reaching in VR, ataxia preserves the visuomotor feedforward and feedback control structure compared to the control group. However, the ataxia group exhibits a small increase in feedback delay (~ 20 ms) and a substantial increase in feedforward delay (~ 70 ms) together with a reduced feedback gain (~25 % lower). Our results suggest that the feedforward and feedback pathways remain largely intact in ataxia, but that time delay deficits and temoral discoordination amongst these control pathways may contribute to the disorder. We also find that providing a preview—analogous to driving on a clear night and seeing the road ahead vs. driving in the fog— improves tracking performance in the ataxia group, although the control group was significantly better able to exploit this preview information. Overall, our results indicate that the feedforward control and preview utilization are relatively well-preserved in individuals with cerebellar ataxia, and that preview could potentially be leveraged to enhance the feedforward performance of those with ataxia.

## 2 Introduction

The study of how animals achieve smooth, consistent movements has long been a topic of interest in biology and robotics. Studies of human movement often focus on the control mechanisms for coordination of limb movements [1], and how damage to different brain areas affect these processes [2, 3]. For example, damage to the cerebellum can lead to incoordination of movement (i.e., ataxia) which results in misdirected and oscillatory reaching, poor balance, and staggering locomotor patterns. A leading hypothesis is that the cerebellum is normally involved in predictive control of movement through the implementation of one or more internal models of the body and environment [4, 5]. Importantly, people with cerebellar ataxia are also unable to effectively adapt their movement to new predictable demands, which is interpreted as an inability to learn and update internal models [6, 7]. As a result, rehabilitation training is often ineffective in improving the movements of individuals with cerebellar damage. Instead, these individuals typically rely on compensatory devices or assistance to reach, balance, and walk.

Previous studies have demonstrated that individuals with ataxia can generate robust online corrections to perturbed feedback, similar to neurologically intact individuals, in various motor tasks, including reaching and speech [6, 8, 9]. Using a system-identification approach [10, 11], we recently showed in a single-joint reaching task that people with cerebellar damage had a visual *feed-back* control structure that was comparable to healthy age-matched subjects [8]. However, people with cerebellar damage had longer time delays than age-matched subjects, potentially resulting in oscillatory movements. Despite this time delay, we could still exploit their intact feedback control structure by designing augmented visual feedback that provided additional phase lead, and this augmented feedback could be customized to improve reaching by correcting an individual patient’s tendency to over- or undershoot targets [8]. Thus, we think that an innovative compensatory solution could be achieved through devices with artificial control designed to actively compensate for the negative effects of ataxia.

Coordinated movement cannot be achieved with only augmented *feedback* control (corrective mechanisms). It also requires *feedforward* control (predictive mechanisms) and the appropriate interaction between feedforward and feedback control. There is limited understanding of how cerebellar damage affects the human feedforward control structure. A specific hypothesis is that the cerebellum is important for feedforward control through its contribution to an inverse internal model of the body [4, 5], i.e., the computation of motor commands to realize desired states, independent of any feedback-dependent corrections. For example, studies have provided behavioral evidence of impairment in feedforward control in people with ataxia, such as the misdirection of the early portion of the movement [12, 13]. Additionally, certain neurological evidence suggests the cerebellum potentially encodes the kinematics of reaching movements [14]. These observations are important, though perhaps owing to the complex interplay between feedforward and feedback control, the specific impact of cerebellar damage on the dynamics of the feedforward controller remains unclear.

In this study, we aimed to develop experimentally validated models that capture the dynamics of how the cerebellum contributes to the feedforward control process and the coordination between feedforward and feedback control. To achieve this, we designed a 3D reference tracking and disturbance rejection task in virtual reality (VR) that enables us to decouple the feedforward and feedback control processes. We systematically quantified how damage to the cerebellum alters the structure (or parameters) of both the feedforward and feedback control processes. Note that humans and other animals can develop internal models of both their own body dynamics and objects in the external environment to improve their control performance [15, 16]. To focus on the role of *internal models of the body dynamics*, the exogenous interactive signals (i.e., the reference signal to be tracked and disturbance to be rejected) were designed to be unpredictable sum-of-sines stimuli [15] to prevent the formation of internal models of these signals [15, 17] and adaptation [9].

Our work revealed that the model structure of the feedforward control process remained largely intact in people with cerebellar ataxia, albeit with a significant increase in feedforward time delay compared to neurologically intact individuals. Based on these results we sought a strategy for leveraging the ataxia group’s intact feedforward control to compensate for their increased delay. Previous research has shown that visual preview of a moving target (i.e., a partial future trajectory) can enhance feedforward control and improve tracking behavior in neurologically intact individuals [18–20]. Such preview information is analogous to seeing a stretch of a curving road out in front of your car while driving [21, 22]. However, it is not well understood how the ability to extract useful information from a visual preview is affected by ataxia. We therefore examined whether preview information could indeed be used by the ataxia group to improve control performance and, if so, how the ability to use preview information is impacted by ataxia. Our findings have implications for understanding the cerebellum’s role in feedforward and feedback control and for the design of external assistive devices to improve performance in clinical settings.

## 3 Results

We studied people with and without cerebellar ataxia as they performed a tracking task using a custom ship-to-home virtual reality (VR) system. We asked them to perform a tracking task where their objective was to keep a white cursor, representing their hand position, as close as possible to a green target (i.e., “reference”) moving pseudo-randomly within a 2D plane (the target could move left-right and up-down). Hand motions were not physically constrained to this plane, requiring 3D, multi-joint visuomotor control by the participants. The white cursor’s movement was pseudorandomly jittered relative to the person’s actual hand position. This “disturbance” provided the participant with perturbed visual feedback of the hand (Fig. 1A). Uncorrelated sum-of-sines were employed to generate the pseudo-random motion of both the reference and disturbance, so that participants could not learn to adapt or predict the movement. This essentially forced participants to perform feedforward motion planning and feedback control at each time point. This design, which is commonly used in system identification experiments [10, 11, 23, 24], allowed us to simultaneously differentiate and analyze the contributions of feedforward control (i.e., planned hand response based purely on the target) and feedback control (i.e., hand response based on the perceived error between the cursor and the target) across a broad frequency range. The control diagram in Fig. 1B illustrates the feedforward and feedback pathways.

**Figure 1:**
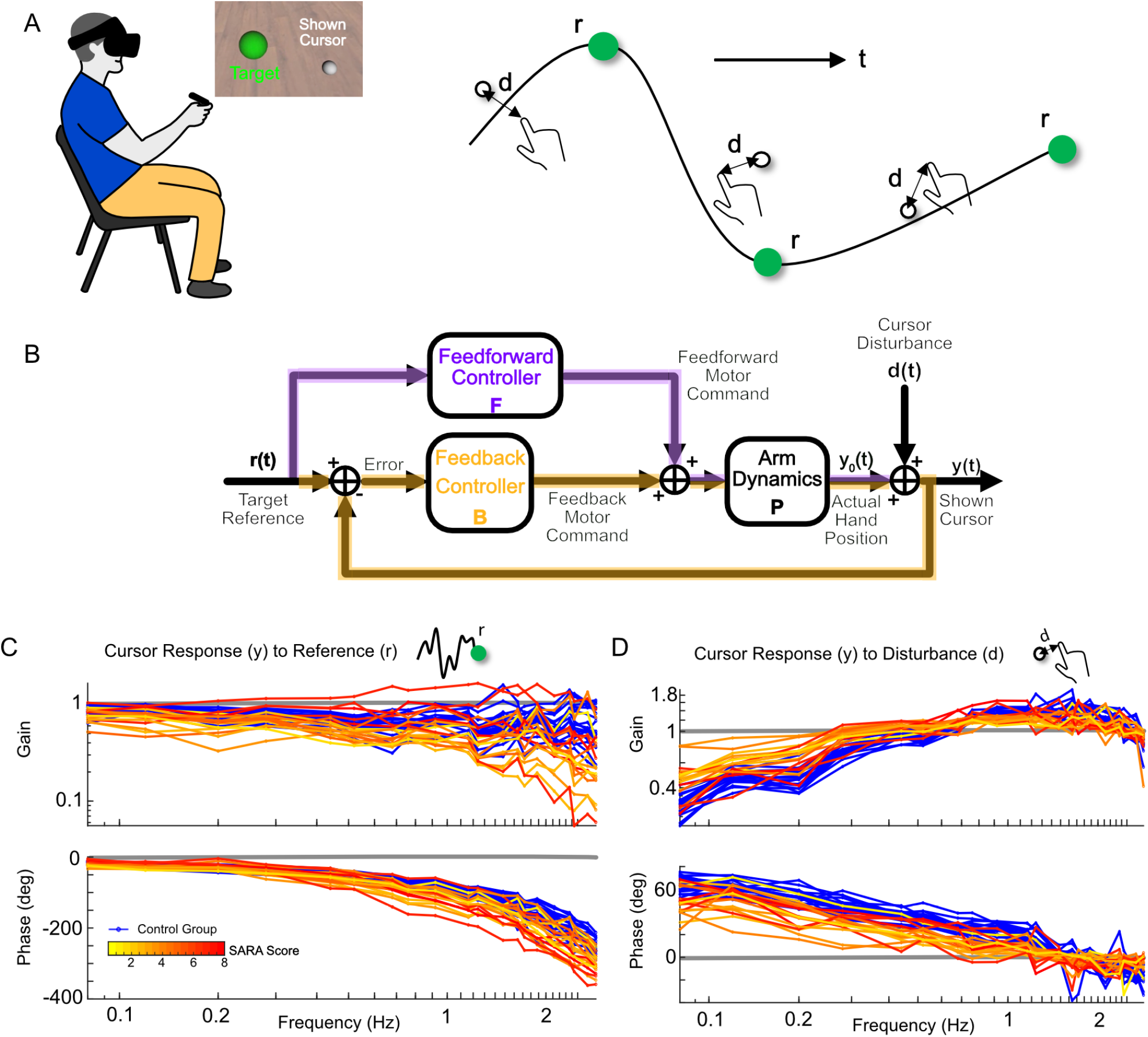
Target tracking design to simultaneously identify both feedforward and feedback control. **A.**Illustration of a participant performing the tracking task with a cropped screenshot of VR display. Participants wore an Oculus Rift S VR headset and held the hand controller which measured the real-time 3D position of the hand. Participants tracked a moving target (green), while their displayed hand position (white) was perturbed by disturbance (d). The target trajectory (r) and disturbance (d) applied to the hand position were uncorrelated, 2D, pseudo-random signals. **B**. Feedback control diagram capturing the experimental topology of the tracking task. The experimental inputs are the location of the visual target reference *r*(*t*) and a disturbance *d*(*t*) that offsets the location of the actual hand position *y*_0_(*t*) before displaying it as a cursor *y*(*t*) to the participant. In this experimental topology, there are two control pathways: feedforward (purple) and feedback (green). The feedforward controller is based purely on the reference, while the feedback controller utilizes the error between reference and shown cursor to generate appropriate adjustments to the motor commands. The combined feedforward and feedback motor commands are fed into the arm dynamics which generate hand movement. **C**. Frequency responses from reference *r*(*t*) to the shown hand cursor *y*(*t*). The control group’s responses are displayed in blue, while the ataxia group’s responses are color-coded based on their upper arm SARA score, which reflects their level of ataxia severity. Control group generally exhibited better tracking (gain closer to 1 and less phase lag) across frequencies than ataxia group (see text). **D**. Frequency responses from the disturbance *d*(*t*) to the shown hand cursor *y*(*t*) revealed that both groups were able to reject disturbances, but ataxia group were able to do so less effectively than control group (see text).

We examined how the designed reference and disturbance signals related to the participant’s hand movements (Fig. 1C,D). A sample of 17 individuals with cerebellar ataxia (ataxia group) and 18 age-matched controls without neurological damage (control group) performed the tracking task. Both groups did well at following the low-frequency components of the reference but experienced increased lag at higher frequencies, with greater variability observed in the ataxia group (Fig. 1C). Both groups were also good at rejecting the disturbance at low frequencies, with the control group demonstrating better rejection, i.e., lower gain to the disturbance (Fig. 1D). Neither group coped well with disturbances at higher frequencies. Individuals with ataxia also exhibited more phase lag in their responses to both reference tracking and disturbance rejection (Fig. 1C,D). Using these responses and following the control diagram, we estimated the feedforward and feedback pathways via frequency response functions (Fig. 2A,B).

**Figure 2:**
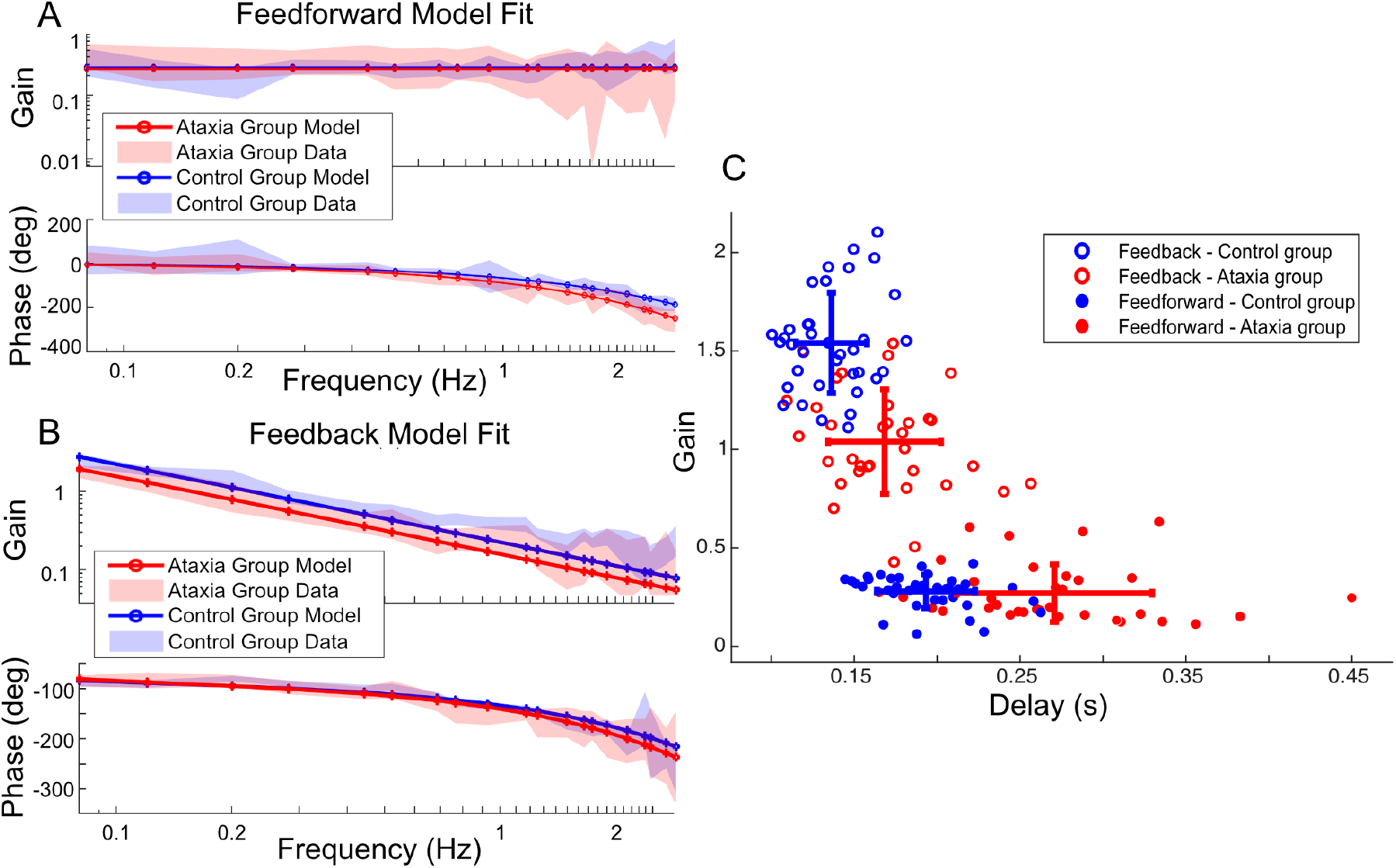
Ataxia group preserves the control structure but exhibits an increase in both feedback and feedforward delays, as well as reduced feedback gain. **A.**Magnitude and phase plots of frequency responses for the feedforward pathway (shaded regions) with a best model fit of the group data (solid lines). The magnitudes are similar across groups but the ataxia group exhibits greater phase lag. The best fit model for both groups is a pure-gain-with-time-delay model, i.e. *ke*^−*sτ*^. **B**. Magnitude and phase plots of frequency responses for the feedback pathway (shaded regions) with a best model fit of the group data (solid lines). The ataxia group exhibits both a lower gain (overall downward shift of Gain plot) and slightly greater phase lag (greater roll off in the Phase plot). The best fit model for both groups is a leaky-integrator-with-time-delay model, i.e. 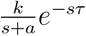. **C**. The parameters (*k, τ*) of the fitted model *ke*^−*sτ*^ for feedforward pathway for all participants. And two of the parameters (*k, τ*) of the fitted model 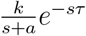 for feedback pathway for all participants. For feedforward pathway, ataxia group show significantly greater time delay than control group (unpaired t test: t(48.01) = 6.88, p < .001, CI = [0.055,0.10]). There was no significant difference between the gain term *k* between groups (unpaired t test: t(52.51) = −0.31, p = 0.79, CI = [−0.067,0.049]). For feedback pathway, ataxia group distributions show significantly smaller gain (ANOVA: F(1,66) = 14.21, p < .001) and significantly greater time delay (ANOVA: F(1,66) = 28.77, p < .001) than control group. The integrator pole *a* (not shown) had no significant group difference (ANOVA: F(1,63) = 0.018, p = 0.89 > .05).

### Ataxia preserves feedforward control, but with greater time delay

We investigated how feedforward control (i.e., inverse internal model control) becomes impaired with the severity of ataxia. In the field of control systems engineering, the feedforward controller is ideally designed to invert the dynamics of the motor plant (i.e., the mapping from muscle commands to movement). Thus it can also be called an inverse internal model—basically, a tool that works backward, effectively computing the requisite motor commands (e.g., the muscle activation) to achieve a desired movement or trajectory based on a reference signal (in our case, the visual target motion); see the purple pathway in Fig. 1B. In this framework, any discrepancies from an imperfect inversion of the plant model are then corrected by feedback; see green pathway in Fig. 1B.

In an idealized setting, the feedforward (inverse internal model) controller would exactly cancel the plant dynamics, and the output would exactly replicate the reference signal. However, real systems have limitations, and perfect inversions are not achievable. For example, since the plant includes sensory and motor delays, perfect inversion would be non-causal. Using system identification techniques, we empirically identified the feedforward control pathway, which is the cascade of the feedforward controller and the plant dynamics. We then described it in terms of how well the feedforward controller can invert the plant. It is in this narrowly defined sense that we claim the brain implements an “inverse internal model” in the feedforward pathway.

We hypothesized that the feedforward control pathway in the control group would be a gain with some time delay. In other words, control participants would be able to achieve the desired feedforward movement, but with a lower gain and some time delay. Damage to the cerebellum may lead to a dysfunctional inverse internal model of the plant dynamics. This could lead to an incorrect version of the feedforward controller that cannot account for all of the plant dynamics. If this were the case, the feedforward control pathway would also have the unaccounted dynamics in addition to the gain with delay. To investigate this hypothesis, we proposed a wide range of candidate models of the feedforward control pathway, such as a pure gain with a time delay (i.e., inverse internal model matches plant dynamics), a first-order model with a time delay (i.e., inverse internal model fails to cancel one mode of the plant dynamics) second-order model (i.e., inverse internal model fails to cancel two modes), etc. The best-fit model for explaining the feedforward frequency responses was selected using a cross-validation process, taking into account the trade-off between bias and variance. More details about the model fitting and selection procedure are described in Methods.

For healthy control participants, we found that a pure gain with time delay model i.e. *ke*^−*sτ*^, where *k* is the gain and *τ* is the amount of delay, was an accurate representation of the feedforward control pathway, as shown in Fig. 2A. This is consistent with previous literature [11] which suggests that the healthy control group may be able to utilize an inverse internal model of the plant in their feedforward controller, with a time delay that arises from sensorimotor and computational delays. We hypothesized that the ataxia group would exhibit a different internal model structure for the feedforward controller (e.g., their best-fit model would be a first or second-order model suggesting a mismatch of the inverse internal model). Instead, we found that a pure gain with time delay model was also an accurate and consistent representation of the feedforward control pathway in the ataxia group, as shown in the Fig. 2A, with one notable distinction: the ataxia group exhibited a significantly longer time delay in their feedforward controller compared to control participants (245 vs. 185 ms; p < 0.001), as shown in Fig. 2B. The significant distinction between the ataxia group and the control group lies in the observed greater phase lag, which can be attributed to the increased time delay. This time delay explanation has a straightforward biological interpretation: it takes longer for the damaged cerebellum to compute or execute the feedforward motion plan. Our results suggest that the impact of cerebellar damage on feedforward control is primarily related to a timing deficit rather than a substantial alteration of the inverse internal model in the feedfoward pathway. See Supplementary Results for additional model-fitting.

### Ataxia preserves feedback control, but with lower gain and greater time delay

We also investigated the relationship between feedback control and the severity of cerebellar ataxia. Based on a previous study examining a single joint task that used reference tracking [8], we hypothesized that the ataxia group would not show a change in their feedback control structure, but would show differences in time delay and gain parameters. We obtained the experimental frequency responses of the feedback control pathway for both groups from the tracking task. We performed model fitting and model selection of the feedback control pathway using different candidate models. The best-fit model for explaining the feedback frequency responses was selected using a cross-validation process, taking into account the trade-off between bias and variance (see Methods).

Our analysis revealed that a leaky integrator with a gain and time delay, i.e.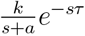, where *k* is the gain, *τ* is the amount of delay, and *a* is the pole of the integrator, accurately and consistently represented the feedback control pathway for both groups (Fig. 2B). However, the ataxia group exhibited significantly greater time delay (144 ms vs. 123 ms; p < 0.001) and significantly smaller gain (1.0 vs. 1.3; p < 0.001) compared to the healthy controls (Fig. 2C). These findings indicate that ataxia led to an increased time delay in the feedback control pathway along with a smaller gain but that otherwise the characteristics of the feedback control pathway were preserved.

### Ataxia increased time delay disparity between feedforward and feedback pathways

As described above, the ataxia group exhibits greater feedforward and feedback delays which adversely affect tracking performance. It is also important that feedforward and feedback control should be coordinated in time. Here we tested how ataxia affects this coordination. Figure 2C shows that people with ataxia had a larger increase in the feedforward time delay than feedback time delay. Figure 3A shows that this difference was significant. The greater time delay difference in the ataxia group suggests compromised coordination between these essential control components, which may also degrade tracking performance. As shown in Figure 3B, we observed a significant, positive correlation between time delay difference and tracking error in the control group. A discernible positive trend was observed in the ataxia group, though this did not reach significance (p = 0.08 > .05).

**Figure 3:**
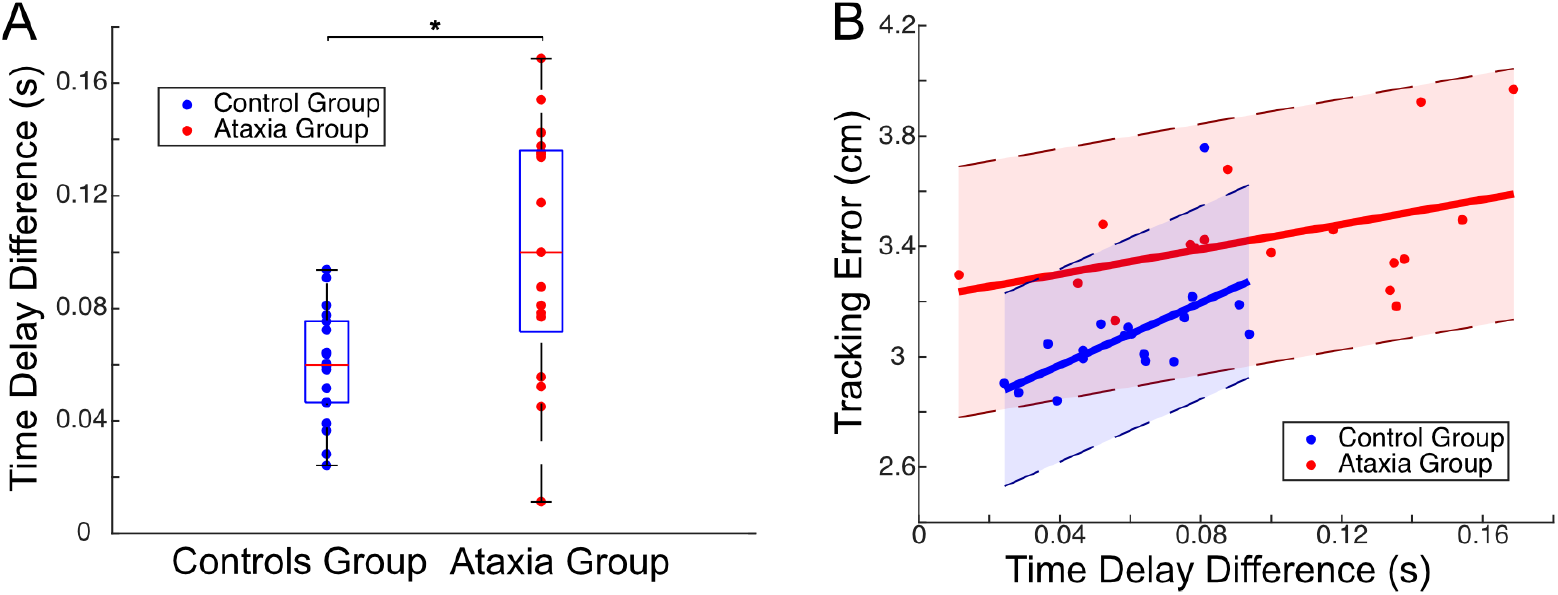
Ataxia induces increased time delay disparity between feedforward and feedback pathways. **A.**Ataxia group shows significantly greater time delay difference between feed-forward and feedback pathway (*τ*_feedforward_ − τ_feedback_) than control group (unpaired t test: t(22.12) = −3.51, p = 0.002 < .05, CI = [−0.066, −0.017]). **B**. In the controls group, the tracking error was found to be positively correlated with time delay difference (r(17) = 0.58, p = 0.01 < .05, 95% CI = [0.16, 0.82]). A discernible (but non-significant) positive trend was observed in the ataxia group (r(16) = 0.43, p = 0.08 > .05, 95% CI = [−0.061, 0.76]).

### Lower feedback gain with ataxia may preserve stability and performance

Above, we showed that the ataxia group exhibits a greater feedforward delay and a greater feedback delay, and these delays are *differentially* impacted which may lead to a discoordination of the feedforward and feedback controllers. A third noteworthy distinction between groups pertains to a reduced gain in feedback control within the ataxia group. If this reduction were caused by an ataxia-induced deficit, we would predict that augmenting the feedback gain to match the control group’s level might recover tracking performance in individuals with ataxia. We simulated this scenario to test this hypothesis. Specifically, we retained all parameters for the ataxia group except for aligning the feedback gain with that of the control group. This did not lead to an improvement in the ataxia group’s tracking; surprisingly, instead it showed a marginal worsening with a slightly higher tracking error, as depicted in Fig. 4A. For more details on how we conducted the simulation, see Methods.

**Figure 4:**
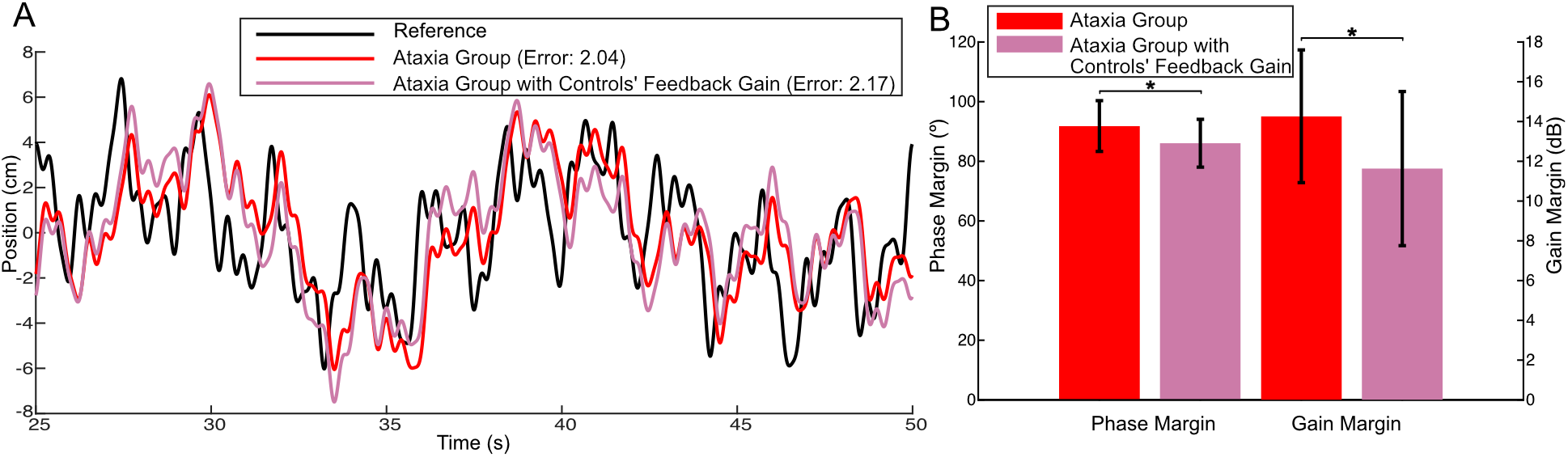
Lower feedback gain with ataxia may preserve stability and performance. **A.**Tracking Simulation. Retaining all parameters for the ataxia group except for aligning the feedback gain with the control group’s level (the pink color) did not lead to an improvement to the ataxia group’s tracking (red); instead, it led to a slightly higher tracking error. **B**. Stability margins (larger is more stable). For ataxia group, aligning the feedback gain with the controls group’s level (the pink color) could lead to a decreased stability margin, in both phase margin and gain margin (unpaired t test: for phase margin, t(65.76) = 2.87, p = 0.006 < .05; for gain margin, t(64.10) = 2.98, p = 0.004 < .05).

In control systems, stability robustness is a crucial property to ensure performance and avoid instability in the face of uncertainty in the feedback system. The *stability margin* quantifies the system’s stability robustness by measuring the distance between the current operating point and the critical point where the system would become unstable (e.g., diverge to infinity). There are different types of stability margins, the most commonly used ones being gain margin and phase margin, as shown in Fig.4B. In both cases, a larger stability margin indicates a more robustly stable system. Notably we found that the reduction in gain among individuals with ataxia contributed to an *increased* stability margin, enhancing their stability robustness, perhaps to compensate for the increased time delay in their feedback controllers (Fig.4B). For details on stability margin calculations, see Methods.

These findings suggest that the diminished gain in feedback control may not be an ataxia-induced deficit. Instead, it could potentially serve as a compensatory mechanism for coping with time delay, thereby conferring a benefit by minimizing the deleterious impacts of the increased delay. We elaborate on this further in the Discussion. It is noteworthy that a similar pattern of gain reduction as a function of increased delay was not observed in the feedforward control pathway. This difference may be attributable to the fact that delay in the feedforward control pathway does *not* impact closed-loop stability robustness and, therefore, there is no benefit to further reduce the gain in this pathway.

### Target preview improves tracking performance, even with ataxia

When driving a car, one is able to see a coming curve in the road before having to make steering adjustments; this is referred to as “preview information,” and can greatly improve tracking performance [25]. We investigated how ataxia, which causes different delays in feedforward and feedback control pathways, could affect the ability to use preview information, and whether such preview information could enhance tracking performance. Our modeling results above indicated that cerebellar ataxia leads to significantly greater time delay in feedforward control, but that the control structure itself remains largely intact. Based on this finding, we hypothesized that a trajectory preview would alleviate tracking deficiencies by providing additional time-lead information about the reference that could be used for feedforward control.

During the preview tracking task, participants tracked a green target while a preview of 500ms of the future target trajectory was shown in blue (Fig. 5A). To test the effect of preview, we performed randomized trials of preview and no-preview tracking tasks as described in Methods. Inspection of representative trials of a control participant (Fig. 5B) and individual with ataxia (Fig. 5C) suggests that the preview information significantly reduces phase lag of the hand trajectory relative to the target for both groups which may reduce tracking error. To test this, we compared the mean squared tracking error in these two conditions for both the ataxia and control groups (Fig. 5D). Both groups showed significant reductions in tracking error with the use of the preview (*p* < .05). These results suggest that *all* individuals were able to effectively utilize preview information. However, the ataxia group exhibited significantly less improvement in tracking error compared to control group (*p* < .05). This is noteworthy as the control group had lower initial tracking error than ataxia group but were nevertheless able to reduce it to a greater extent. This demonstrates that ataxia impacts how effectively preview information is utilized.

**Figure 5:**
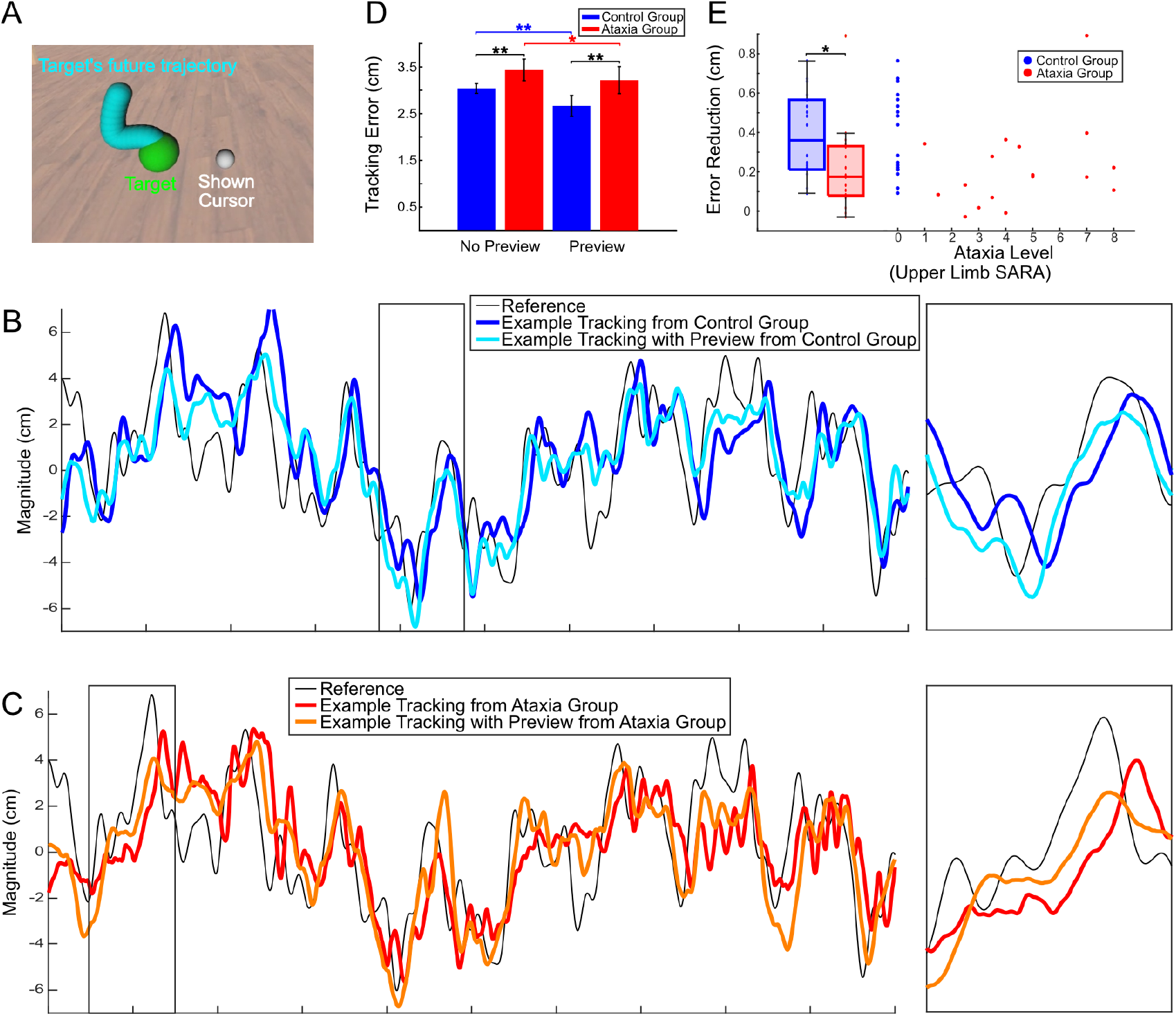
Both populations improve tracking performance with preview information, but ataxia group exhibited significantly less improvement. **A.**During preview tracking, participants were shown a 500ms preview (blue) of the target’s future trajectory. **B**. Hand tracking trajectories with (cyan) and without (blue) target preview from an example control participant tracking a reference trajectory (black). Inset (2 s) shown in right panel. Note that the hand trajectory with preview leads in phase compared to the trajectory without preview. **C**. Hand tracking trajectories from an example participant with ataxia, similar to panel D. Note again that the hand trajectory with preview (orange) leads the hand trajectory without (red), just as with control participants. **D**. Comparison of tracking error between preview and no-preview conditions. A two-way ANOVA (group × condition) was performed with post-hoc analysis and Bonferroni correction. There is a significant difference in tracking error between ataxia group and control group (black lines) for both the no preview condition *(p* < .*001)* and the preview condition *(p* < .*001)*. And the presence of the trajectory preview significantly reduced tracking error for both ataxia group (red line) *(p =* .*034)* and control group (blue line) *(p* < .*001)*. **E**. The ataxia group shows significantly less tracking improvement than control group *(p = 0*.*04* < .*05)*. No significant correlation between SARA Score and the tracking error reduction, as calculated by subtracting the mean tracking error with preview from the mean tracking error without preview*(r(17) = 0*.*25, p =* .*37* > .*05, 95% CI = (−0*.*25 0*.*63))*.

To further analyze the impact of ataxia on preview utilization, we examined if there is a relationship between the level of ataxia and the amount of tracking improvement caused by the presence of preview. We found no significant correlation between the level of ataxia and the error reduction caused by preview (Fig. 5E). This implies that the benefits of path preview are impacted by ataxia, but may not depend directly on the severity of the pathology. These findings have important implications for future rehabilitation and therapeutic approaches, as they demonstrate the potential of a mechanism for performance improvement that can confer modest benefits regardless of the severity of the ataxia.

### Ataxia reduces phase lead and attenuates gain in preview utilization

In addition to examining the impact of preview information on control performance, we also sought to empirically model the ataxia-induced deficits in preview utilization.

The feedforward and feedback control diagram (Fig. 1B) is insufficient to assess preview tracking because subjects can also incorporate other time points within the preview to guide their feedback control. To address this, we used a more comprehensive two-channel model consisting of a reference control pathway based purely on the reference signal including preview information, and a feedback control pathway based on the hand cursor information (see Methods and Fig. 9 for more details about the two-channel model).

To investigate the utilization of preview information by subjects, we compared the cursor response (*y*) to reference (*r*) between the preview condition and non-preview condition (Fig. 6A).

**Figure 6:**
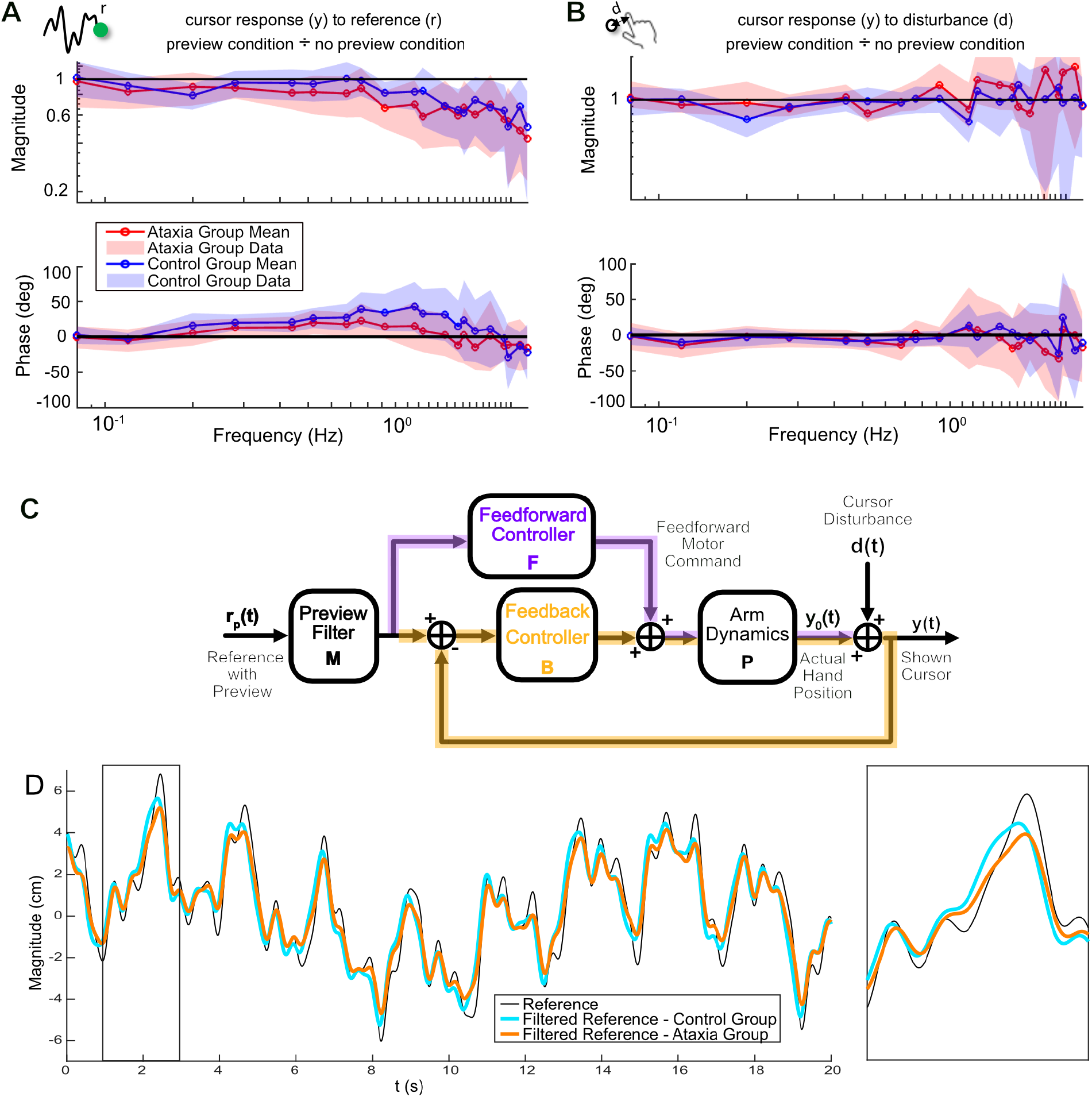
Preview information adds phase lead while filtering out high frequencies in feedforward controller. **A.**Magnitude ratio and phase differences of the *reference control pathway* showing group means ± SD for both ataxia (red) and control (blue) groups. Preview conferred two major benefits for both groups: gains were attenuated at higher frequencies (rolling off below 1, top plot) and significant phase lead was added across a wide bandwidth (phase above 0^°^, bottom plot). **B**. Same plots for *disturbance rejection (feedback) control pathway* shows negligible change in gain or phase, indicates that preview information had little-to-no effect on feedback control as expected. **C**. Control theoretic model for preview tracking that includes a preview filter, *M*, which filters the preview information, *r*_*p*_(*t*). This filtered reference is then processed by the feedforward and feedback controllers precisely as in the no-preview condition (Fig 1B). **D**. The filtered reference represents the target trajectory participants are tracking under the preview condition, adjusted by each group’s preview filter. For both groups, the filtered reference (cyan and orange) is smoother than the original reference signal (black) with higher-frequency details filtered out (e.g., fewer sharp turns), and the filtered reference signal is phased advanced relative to the unfiltered reference (e.g., peaks and reversals in the filtered reference occur earlier).

The results revealed that the preview information provided more phase lead (predictive) in the low frequency range and a gain drop-off at high frequencies for both ataxia group and control group. However, individuals with ataxia showed less phase lead than controls and the gain began dropping off at an earlier frequency. We also assessed the impact of preview information on the feedback control pathway by examining the changes in the cursor response (*y*) to disturbance (*d*) with and without preview (Fig. 6B). We found no appreciable change in the magnitude (ratio is around 1) or phase (difference is around 0 deg) between conditions. These results matched our prediction that the reference preview would not affect the feedback controller

Based on the fact that preview introduced phase lead to the reference utilization, we modified our original feedforward and feedback control scheme by including an additional reference preview filter (Fig. 6C). In this model, the reference signal together with the preview is filtered through the reference preview filter and then sent to the original feedforward and feedback control pathways. More details about how we derived this model are described in Methods. By comparing the preview filters for ataxia group and control group, we can explore the role of the cerebellum in the utilization of preview information. As shown in Fig. 6D, for both groups, the filtered reference is smoother than the reference signal with higher-frequency details filtered out (e.g., fewer sharp turns, better matching the bandwidth of the downstream motor system), and the signal is slightly phase leading compared to the unfiltered reference (e.g., peaks and reversals in the filtered reference occur earlier in time, thanks to the utilization of preview information). These improvements (smoothness and phase lead) were stronger in the control group than in the ataxia group. These models support the finding that ataxia impairs—but does not eliminate—the ability to extract information from the preview trajectory, leading to less effective utilization of this information in the control process.

## 4 Discussion

In this study, we investigated the distinct characteristics of feedforward and feedback control mechanisms in individuals with ataxia compared to a control group during a visuomotor tracking task. Visually guided tracking movements are commonly used in clinical tests to assess the severity of ataxia [26]. Previous studies of people with ataxia have focused on timing and kinematic parameters during tracking movements, but have not dissociated feedforward and feedback control architectures [27–29]. One study addressed whether cerebellar patients show different movement patterns when reaching in the dark versus light, assuming that feedforward control would be dominant in the dark [30]. They found that feedforward hand responses were abnormal in the dark, whereas in the light, oscillatory feedback corrections helped reduce errors. Here we developed models that allow us to quantitatively describe the type of control structure being used for feedforward and feedback mechanisms.

We know of no other work that has developed experimentally validated models to understand cerebellar contributions to feedforward and feedback control. Our approach was to introduce an unpredictable disturbance to the displayed cursor that was uncorrelated to the target motion [10, 11, 31]. We measured visual feedback control (i.e., the hand response to the perceived error between the cursor and the target) by assessing how well participants could reject the cursor disturbance. After accounting for this visual feedback response, we were able to characterize feedforward control (i.e., the planned hand response based purely on the target). In other words, feedforward control during tracking involves continuously predicting where the hand should move given the target position, without relying on the servo-like corrections of the observed cursor to target error.

We estimated internal model structures for feedforward and feedback control, and uncovered significant differences between people with cerebellar ataxia and controls. We were surprised to find that the internal model controller in the feedforward pathway may not be structurally altered by cerebellar ataxia. The major change that was evident from the frequency response function was an increased phase lag, consistent with a 70 ms longer feedforward time delay compared to controls. There was no change in feedforward gain. The structure of the feedback controller was also preserved in the ataxia group, but operated with a 20 ms longer delay and a smaller gain relative to controls. The smaller gain may serve as a conservative means of mitigating effects of feedback delay in patients because it helps to maintain stability robustness. Finally, while we found that both groups were able to benefit from “visual preview” (i.e., the displayed phase lead information), the ataxia group showed less of an improvement.

These findings are significant as it has previously been challenging to quantitatively model feedforward control independently of visual feedback control in ataxia patients. We uncovered different time delays in the feedforward vs. feedback pathways, suggesting that the cerebellum may be important for the temporal coordination of these two pathways. The ability of ataxic individuals to utilize preview information, albeit less somewhat effectively than controls, suggests that enhancing visual compensation strategies could potentially improve motor control in individuals with ataxia.

### Longer Time Delay in Feedforward Control Identified in Ataxia

We found that the feedforward control pathway—comprising the feedforward controller and plant dynamics—functions as a pure gain with some time delay for both the control and ataxia groups. Interestingly, the main difference between the groups lies in the phase, which can be attributed to a longer time delay, and minimal group differences in the gain. This suggests that their internal model controller might not be substantially altered due to cerebellar damage. Instead it simply operates at a much longer delay compared to that of controls.

Previous literature has hypothesized that cerebellar ataxia is due to dysfunctional or impaired internal models [5, 32, 33]. For example, Bhanpuri et al. [33] showed in simulation that the inertial mismatch between the inverse internal model (feedforward controller) and plant dynamics could potentially explain the dysmetria pattern seen in people with ataxia. Our results, however, do not find clear evidence of an internal model mismatch from the plant (i.e., arm dynamics), but may support other possible mechanisms. A novel finding is our observation that damage to the cerebellum results in longer and *different* time delays in feedforward versus feedback control pathways. This may reflect an impairment in temporal coordination and synchronization between these control components, contributing to the movement difficulties observed in individuals with cerebellar ataxia. We plan to explore this possibility further through simulations in future work.

It is well-known that people with cerebellar damage exhibit impaired feedforward control performance, typically characterized by longer delay in movement initiation for eye saccades and hand motion [34–36], and larger initial deviations [12, 13], especially observed in the ballistic movement phase of reaching. Our results add to the literature by demonstrating that the ataxia group exhibits approximately 70 ms longer for the preparatory phase in response to continuous but unpredictable target motion. However, our experimental results on the cerebellar contribution to feedforward control do not provide an explanation for the larger deviation observation. First, note that the initial deviation pattern is typically direction-dependent. And this observation could be potentially explained by the impaired or mismatched compensation for intersegmental dynamics between joint movements [37, 38]. Second, the discrepancy our findings show may be attributed to the fact that our task is designed as a continuous, dynamic movement tracking task as opposed to point-to-point reaches. The feedforward mechanism we identified from the tracking task may represent the average responses across all directions.

Finally, we noted that the feedforward gain is smaller than 1 and not statistically different between both groups in this experiment. We think that the low feedforward gain was due, in part, to the use of a pseudo-random stimulus. The random nature of the stimulus and the disturbance will push subjects to rely on a combination of feedforward and feedback control, with feedback control playing a more prominent role. Thus, it makes sense why this demand would naturally limit the contribution (i.e., gain) of feedforward control.

### Smaller Gain and Longer Delay Characterize Feedback Control in Ataxia

For both the control group and ataxia group, feedback control can be approximated as a scaled leaky integrator with some time delay, although the ataxia group exhibits longer delay with significantly smaller gain. The feedback response in both groups aligns with the classical McRuer Crossover model [39], which captures the sensorimotor frequency response near the gain crossover frequency (i.e., where the open-loop gain of the combined plant and controller has unit magnitude). In other words, our results show that both groups’ visual feedback responses to unexpected visual perturbations are proportional to the accumulated error signal, with some leakage and time delay. Notably, our approach here considers the feedback responses *after accounting for feedforward contributions*, providing a more nuanced evaluation of feedback control compared to our previous study [8].

The ataxia group exhibited a significantly greater time delay compared to the control group (144 ms vs. 123 ms; p < 0.001). This finding aligns with previous reports on visual feedback delays, which generally range from 110 to 160 ms [40–45] and further validates our prior work showing delayed feedback mechanisms in individuals with ataxia [8]. We suspect that the smaller gain observed in the ataxia group may not be a deficit induced by cerebellar damage. Instead, the smaller gain contributes to improved tracking performance and enhanced stability robustness in the face of a longer time delay. That being said, a smaller gain at longer delay can still lead to the oscillations typically observed in the ataxia group [46]. In other words, while the smaller gain acts to reduce the amplitude of corrective movements in ataxia, it does not completely mitigate them due to the longer time delay. Therefore, people with ataxia can still make larger corrections than control subjects.

An interesting new finding is that the time delays in feedforward and feedback control are inappropriately coordinated in time in the ataxia group relative to controls. While we do not yet understand the full ramifications of this finding, our work here suggests that poor temporal coordination of these two mechanisms may contribute to the movement incoordination in people with ataxia. In the next section, we describe how our manipulation of visual preview information may help normalize inappropriately coordinated timing between feedforward and feedback control mechanisms. Our future work will investigate ways to manipulate timing of visual and proprioceptive feedback and determine if this can reduce ataxia.

### Preview Enhances Tracking in Ataxia and Control Groups

Lastly, we investigated the role of preview information in improving tracking performance in both the ataxia group and the control group. Critically, our results demonstrate that both groups are able to make use of preview information, although the improvement was reduced in the ataxia group. To understand possible reasons for this finding, we modeled the effects of preview tracking and explain a potential cerebellar contribution to preview utilization.

We modeled preview tracking by implementing a preview filter and feeding the predicted target information into the feedforward and feedback control scheme [19, 20]. As shown in Fig. 6 and Supplementary Information, preview utilization is not a simple lowpass filter. Instead, individuals tend to extract the low-frequency components of the future information, while also maintaining the high-frequency component near the target (i.e., the tail of the preview information). The ataxia group extracted less phase lead information from preview and attenuated the gain magnitude compared to controls. Despite these deficits, the ataxia group was still able to utilize preview control. The preview information provides a signal that is similar to use of high-beam headlights when driving on a dark, winding road. People can see farther along their desired path, and this allows them more time to plan a feedforward motor command.

Preview information is easy to provide when a person is moving to track an external stimulus or reference, but may be more challenging when there is no target to track. In that case, this compensation would require the ability to display preview of a the movement intention in advance, for example in an augmented reality display. In order to use such an approach for training movement, it may be necessary to restrict the use of preview compensation to a limited number of common daily activities where the preview design can be trained in advance. In the future, it may be feasible to predict movement intentions using physiological signals reflecting neural intention (e.g., electromyography).

## ACKNOWLEDGMENT

This work is supported by the National Science Foundation (1825489 and 1825931), National Institutes of Health (R01-HD040289), Kavli Neuroscience Discovery Institute (Distinguished Graduate Fellows 2022).

## 5 Methods

### 5.1 Participants

In this study, we recruited a sample of 17 individuals with cerebellar ataxia (ataxia group) and 18 age-matched healthy controls (control group). Individuals with cerebellar ataxia were excluded if they had any clinical evidence of damage to extra-cerebellar brain structures, or clinical evidence of dementia, aphasia, peripheral vestibular loss, or sensory neuropathy. The age-matched controls were also screened for any neurological impairments. Ataxia severity was quantified using the Scale for Assessment and Rating of Ataxia (SARA) score [26]. Since this task is an upper limb tracking task, we calculated an Upper Limb (UL) SARA sub-score comprising the sum of the upper-limb categories of the SARA test, including the finger chase, nose-finger test, and fast alternating hand movements. This clinical test was performed by Dr. Amy Bastian.

All participants gave informed consent according to the Declaration of Helsinki, and the experimental protocols were approved by the Institutional Review Board at Johns Hopkins University School of Medicine. Each participant used their dominant arm during the tracking task. Demographic and clinical information for the participants with ataxia is presented in Table 1.

**Table 1:**
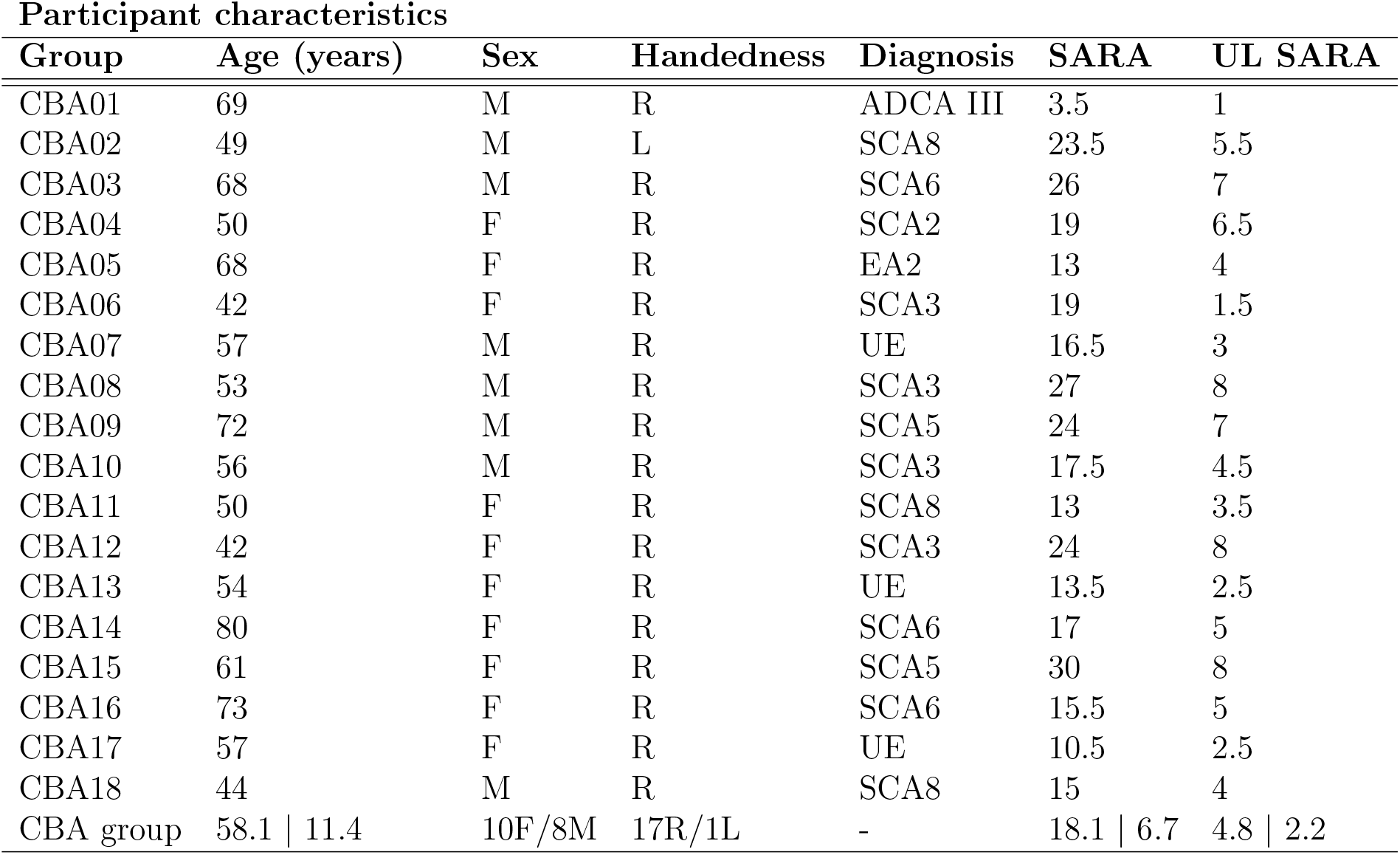
Participant demographic and clinical information.

| Group | Age (years) | Sex | Handedness | Diagnosis | SARA | UL SARA |
| --- | --- | --- | --- | --- | --- | --- |
| CBA01 | 69 | M | R | ADCA III | 3.5 | 1 |
| CBA02 | 49 | M | L | SCA8 | 23.5 | 5.5 |
| CBA03 | 68 | M | R | SCA6 | 26 | 7 |
| CBA04 | 50 | F | R | SCA2 | 19 | 6.5 |
| CBA05 | 68 | F | R | EA2 | 13 | 4 |
| CBA06 | 42 | F | R | SCA3 | 19 | 1.5 |
| CBA07 | 57 | M | R | UE | 16.5 | 3 |
| CBA08 | 53 | M | R | SCA3 | 27 | 8 |
| CBA09 | 72 | M | R | SCA5 | 24 | 7 |
| CBA10 | 56 | M | R | SCA3 | 17.5 | 4.5 |
| CBA11 | 50 | F | R | SCA8 | 13 | 3.5 |
| CBA12 | 42 | F | R | SCA3 | 24 | 8 |
| CBA13 | 54 | F | R | UE | 13.5 | 2.5 |
| CBA14 | 80 | F | R | SCA6 | 17 | 5 |
| CBA15 | 61 | F | R | SCA5 | 30 | 8 |
| CBA16 | 73 | F | R | SCA6 | 15.5 | 5 |
| CBA17 | 57 | F | R | UE | 10.5 | 2.5 |
| CBA18 | 44 | M | R | SCA8 | 15 | 4 |
| CBA group | 58.1 \| 11.4 | 10F/8M | 17R/1L | - | 18.1 \| 6.7 | 4.8 \| 2.2 |

### 5.2 Experimental Design and Procedure

Due to the COVID-19 pandemic, we developed a custom “ship-to-home” virtual reality (VR) system for remote data collection. The system includes a high-performance laptop equipped with Intel Core i7, 32GB RAM, 1TB NVMe SSD + 2TB HDD, NVIDIA GeForce RTX 2080 8GB, 15.6” FHD IPS 144Hz 3ms, Windows 10, and an Oculus Rift S VR headset. The setup was shipped to the participants’ homes for use in the study. Investigators interacted with participants via Zoom and ran experiments via remote access to the laptop. Importantly, an investigator was present on Zoom to monitor each participant as they performed the tasks described here. Participants sat in a desk chair with back support but without wheels or arms and used the VR headset and hand controller to perform the VR tracking task (Fig. 7A). The real-time 3D hand position was measured through the hand controller and rendered in the VR scene at 80 Hz. The VR task was completed via the local computer software and not via internet to ensure there was no internet-connection-based delays in the tracking and data collection.

**Figure 7:**
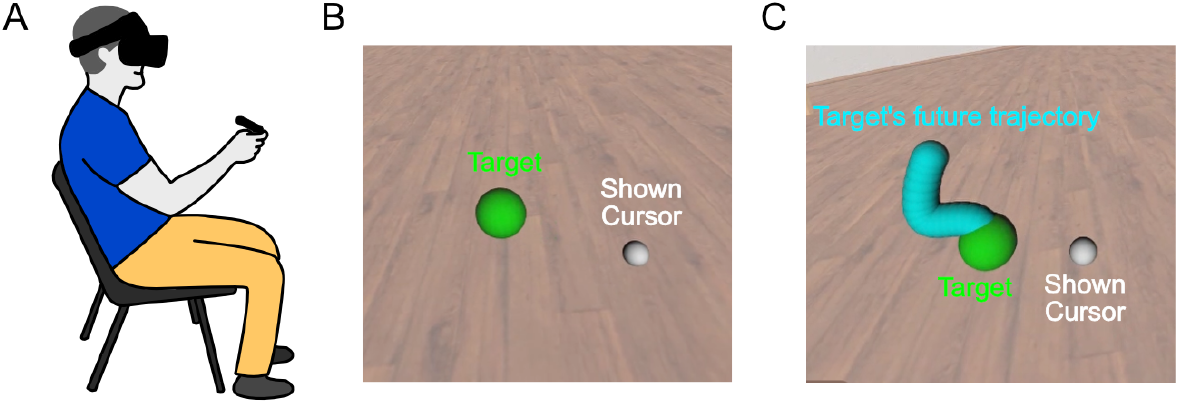
Custom VR system to simultaneously identify both feedforward and feedback control during target tracking. **A.**Illustration of a human performing the VR task. Participants wore an Oculus Rift S VR headset and held the hand controller which measured the real-time 3D position and orientation of the hand. **B**. Cropped screenshot of VR display (no preview tracking). Participants tracked a moving target (green), while their hand position (white) was perturbed. The target trajectory and disturbance applied to the hand position were uncorrelated, 2D, pseudorandom signals. **C**. Cropped screenshot of VR display (with-preview tracking). In the preview tracking, participants were also shown a 500ms preview of the target’s future trajectory (blue).

We conducted two tracking tasks, one without preview and the other with preview. In no-preview tracking (Fig. 7B) participants tracked a 2D, pseudo-random reference (green target) in a 3D virtual reality (VR) environment. Meanwhile, a second uncorrelated pseudo-random 2D disturbance was applied directly to the participant’s actual hand position, and the perturbed hand position (white cursor) was displayed in VR. In the with-preview tracking task (Fig. 7C), eparticipants were also given a 500ms preview of the future trajectory (blue) of the green target.

We used the feedforward and feedback control paradigm depicted in Fig. 1B. Using two input channels—reference (*r*) and the disturbance (*d*)—reveals the contributions of feedforward control (i.e., hand response based purely on the target) and feedback control (i.e., hand response to the perceived error) simultaneously across a broad frequency range [11, 23, 24]. By examining the relationship between input signals and participant’s motion (hand position) using a control-theoretic framework, we quantitatively characterized the feedforward and feedback pathways through frequency response functions.

The target reference and disturbance signals were designed to be unpredictable (pseudorandom) sum-of-sines stimuli [15] to ensure the participants could not form internal models [17] of the exogenous signals to improve their performance; rather, the use of unpredictable stimuli focuses our attention on the role of internal models of the human body-dynamics in tracking [5, 6, 16]. The two input channels in a 2D plane require four sets of sum-of-sines signals, one for each direction (x and y axis) per channel (reference and disturbance). We selected the first twenty prime multiples of 0.04 Hz, ranging from 0.08Hz to 2.84Hz, to form the four sets of sum-of-sines signals, each composed of five unique frequencies. For a given frequency group *g*_*k*_, the sum-of-sines signal was expressed as follows:

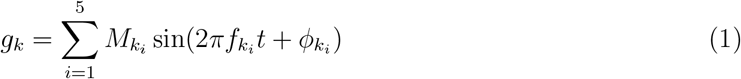

Here, 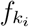 represents a prime multiple of 0.04 Hz, 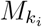 is magnitude of the sine signal, 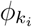 is a random number phase shift, and each group *k* has a different set of 5 sine signals (i.e. a different set of prime multiple frequencies). The magnitudes of the sine signals were scaled to remain within both velocity and positional limtis, similar to [47]:

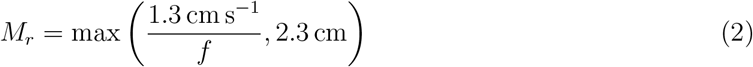

This magnitude profile ensures that *V*_*r*_ ≤ 2.6π cm s^−1^ and *M*_*r*_ ≤ 2.3 cm. To prevent the participants from being overly distracted by the disturbance signal, the velocity and magnitude boundary of the disturbance signal are kept smaller than that of the reference as follows:

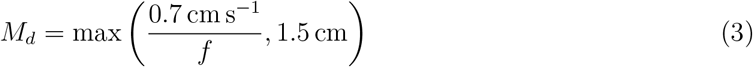

The frequency composition of these groups and a time domain example of a signal group can be seen in Fig. 8.

**Figure 8:**
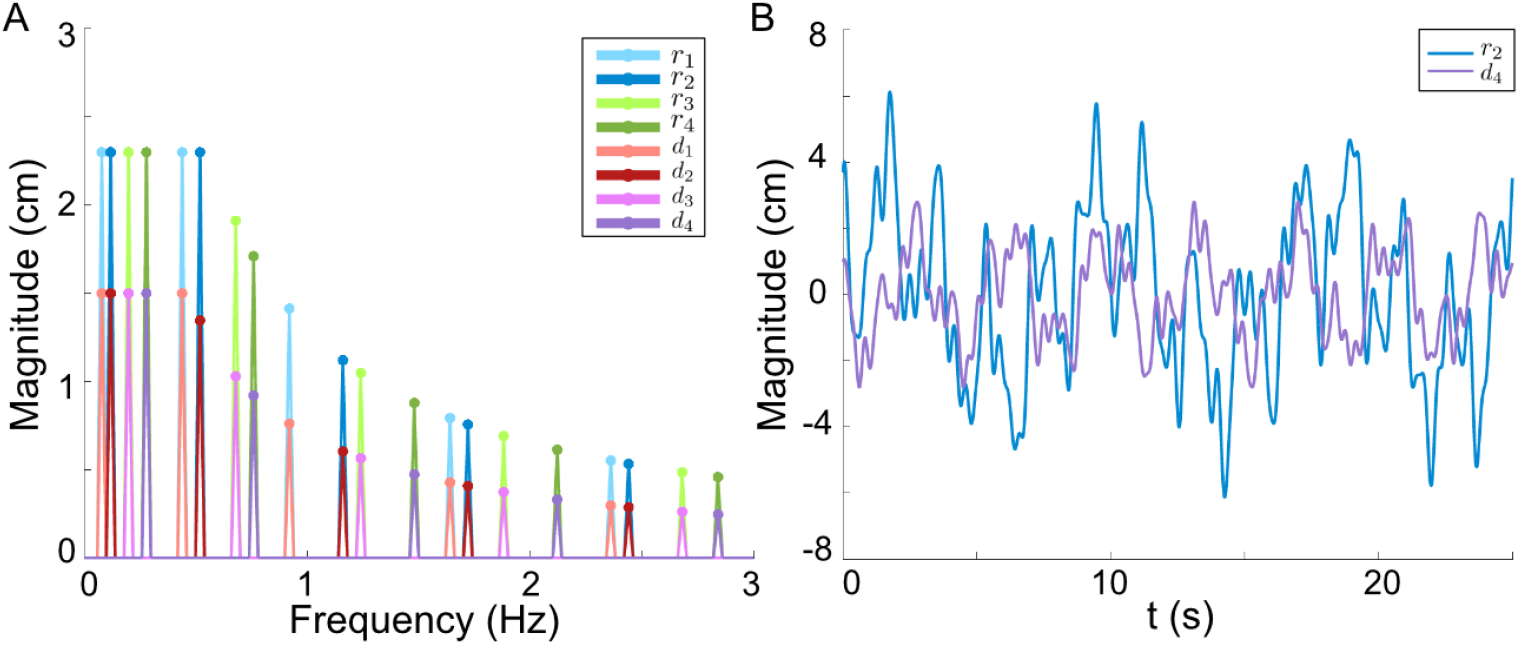
Reference and Disturbance Signals. **A**: Frequency composition of the sum-of-sines groups for reference (*r*_1_, *r*_2_, *r*_3_, *r*_4_) and disturbance (*d*_1_, *d*_2_, *d*_3_, *d*_4_). **B**: Example time-domain traces of a sum-of-sine reference (*r*_2_) and disturbance (*d*_4_).

**Figure 9:**
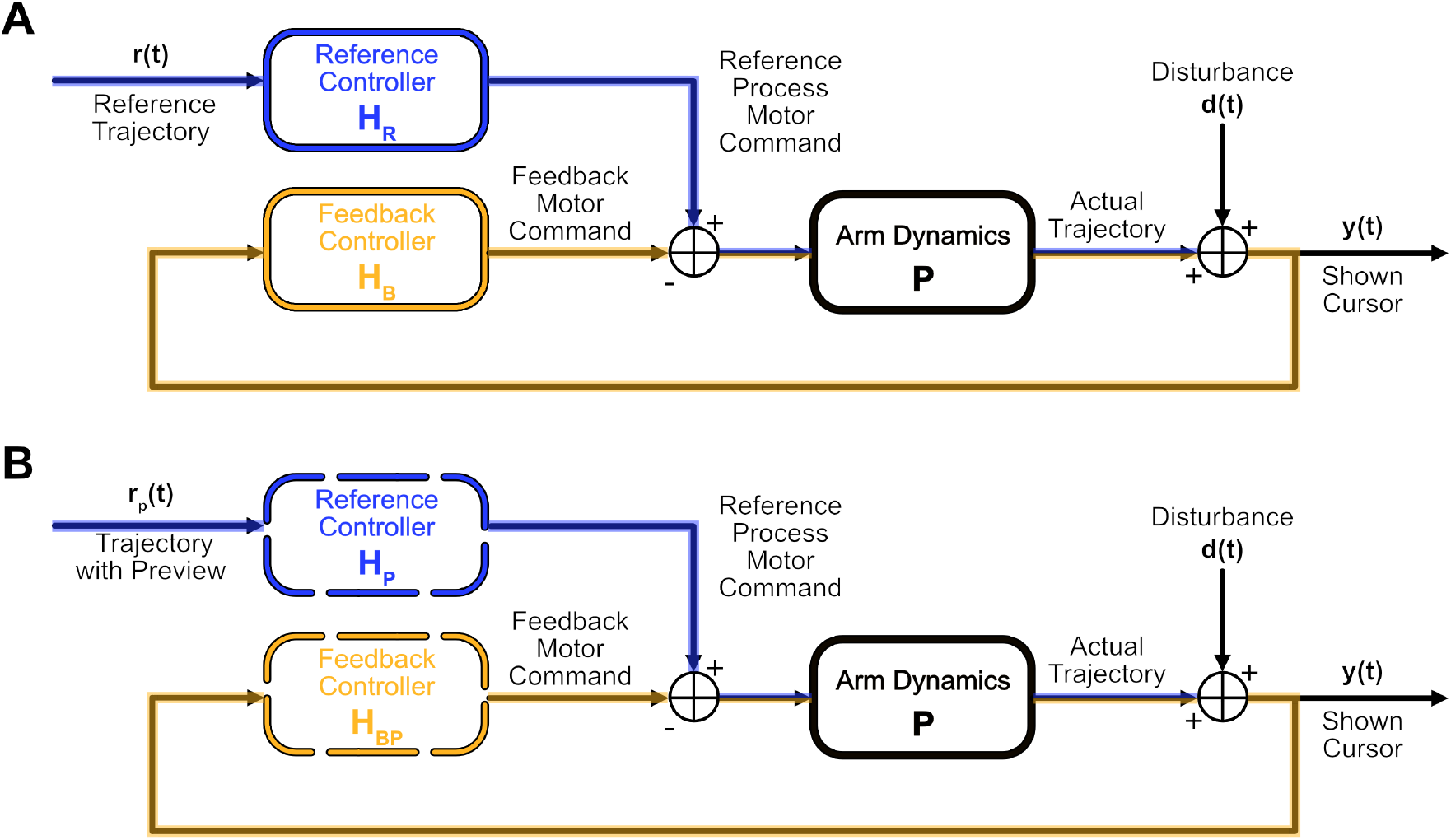
Reference-controller based model. **A.**No-preview tracking model. *r*(*t*) is reference signal at a given time-point; *d*(*t*) is disturbance signal and *y*(*t*) is shown perturbed hand position. *H*_*R*_ is the controller that processes the reference then puts the processed motor command through a feedback loop including the arm dynamics. **B**. Preview tracking model. *r*_*p*_(*t*) is reference signal that includes all time-points in the 500ms preview, *d*(*t*) and *y*(*t*) are the same as no-preview condition. *H*_*P*_ is the controller that processes the full reference signal with preview to generate a motor command, then puts the processed motor command through the same feedback loop as no-preview condition.

Each set of sine signals *g*_*k*_ is then assigned to *r*_*x*_, *r*_*y*_, *d*_*x*_, and, *d*_*y*_ respectively for a given trial. For the trials, sets of sum-of-sines signals are implemented under switched conditions between reference and disturbance and switched conditions for the x and y axis (the two axis for 2D plane). This ensures we have input–output data of all frequency components for the reference and disturbance, as well as all frequency components for each axis of motion. Though the sum-of-sines signals are not predictable, we pseudo-randomize which set *g*_*k*_ is assigned to the x and y components of *r* and *d* respectively for each trail.

To investigate how the ability to predict the reference improves performance, we have two tracking conditions: one with no target preview and one with a 500ms target preview. Each subject undergoes 20 trials: 4 preliminary practice trials, 2 with preview and 2 without, and 16 experimental trials, 8 with preview and 8 without preview. The preview and non-preview trials are also pseudorandomly assigned and interspersed to ensure non-predictability. An example assay of trials and their signal sets can be seen in Table 2.

**Table 2:** Signal composition example of the testing session. Each trial was assigned a frequency composition and preview condition in a pseudo-random manner, to eliminate any potential for predictability. In addition, this assignment was implemented to ensure each frequency component is measured for reference and disturbance in both directions. The bolded numbers indicate specific trial instances. Trials 1 to 4, not presented in the table, were designed primarily for instructive purposes to ensure participants understood the requirements of the tracking task. As a result, these initial four trials were excluded from data analysis.

| Tracking Trial Numbers |  |  |  |  |  |  |  |
| --- | --- | --- | --- | --- | --- | --- | --- |
| Frequency Composition |  |  |  | Preview Time (ms) |  |  |  |
| $r_x$ | $d_x$ | $r_y$ | $d_y$ | 0 | | 500 | |
| $r_1$ | $d_2$ | $r_3$ | $d_4$ | **9** | **13** | **8** | **18** |
| $r_2$ | $d_1$ | $r_4$ | $d_3$ | **5** | **16** | **12** | **19** |
| $r_3$ | $d_4$ | $r_1$ | $d_2$ | **11** | **20** | **6** | **14** |
| $r_4$ | $d_3$ | $r_2$ | $d_1$ | **7** | **17** | **10** | **15** |

### 5.3 Feedforward and Feedback Pathway Isolation

In this experiment, we aim to analyze the feedforward and feedback control pathways involved in the tracking task using only the available information of the designed inputs, reference and disturbance, and recorded output, the hand position. Since we do not have direct access to these pathways, we must estimate them computationally. Utilizing the control diagram (Fig. 1), we can evaluate the system through the following equations (note the (*jω*) notation denotes the Fourier Transformation of the respective time domain signals):

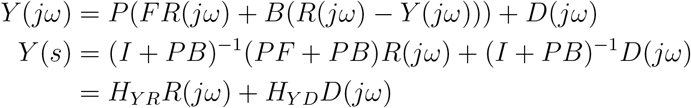

*H*_*Y R*_ represents the frequency response function from output to reference. Likewise *H*_*Y D*_ represents the frequency response function from output to disturbance. By utilizing these frequency response functions, we can calculate the open loop feedforward pathway (*P F*) and open loop feedback pathway (*PB*) as follows:

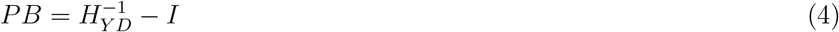

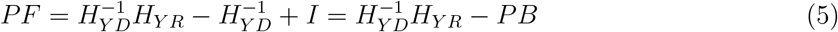

This provides us with a computational approach to determine the frequency responses of the feed-forward and feedback pathways from the behavioral data from the closed-loop control.

### 5.4 Computational Modeling

To understand the control mechanisms of the feedforward and feedback pathways, we aim to fit numerical models to our calculated frequency domain data for the feedforward (*PF*) and feedback (*PB*) pathways. Our objective is to find models that have low fitting error, reasonable dimensionality, and good cross-validation performance without overfitting. We consider various model candidates for both the feedforward pathway and feedback pathway respectively. For instance, one model candidate for the feedback pathway in the frequency domain is the McRuer Crossover model [39] which takes the form 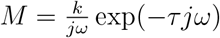, a delayed, scaled integrator. Further information on the model candidates is provided in the Supplementary section.

For each subject, we represented their frequency response as an array of 20 complex numbers, one for each frequency tested in the sum-of-sines experiment, that contain input-to-output gain and phase information at each frequency. The *x* and *y* axis were fitted separately, and there was no significant off axis motion to merit a model fit (see Supplemental Materials). The following procedure was repeated for each possible model candidate. First we would select a candidate model (e.g., McRuer Crossover). Then, for a given group (e.g. the ataxia group) we pulled one subject’s data (e.g. subject *i*) out of the group and took the average of remaining frequency response data in the group at each tested frequency. Using the MATLAB function fminsearch, we then fitted the parameters of the selected model to these average frequency response functions by minimizing the frequency-domain (FD) error (mean squared error):

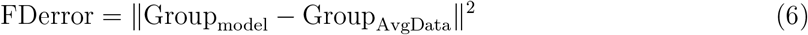

This was repeated 100 times with different initial parameter values selected for the ‘fminsearch’ function, increasing the likelihood of finding a global minimum. The model parameters that generated the lowest fitting error and the magnitude of that error were kept. To verify how well the model generalizes, we then calculate the validation error (VE) between the model fit result and the subject-i data as follows:

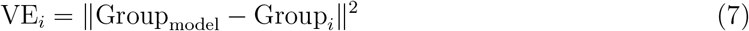

We repeat the above process by leaving out a different subject each iteration, fitting new model parameters, then calculating the VE for each subject. Once we loop through all subjects, the cross-validation error (CVE) is calculated as the average of all these VE errors:

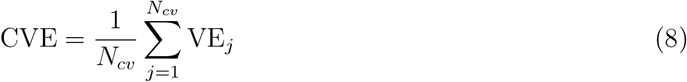

We obtain a CVE and a fitting error for each model candidate. The model candidate that has the smallest CVE and fitting error, while being as simple as possible, is selected.

### 5.5 Preview Filter Extraction

Here we present how we derive the control diagram of preview tracking in the Fig. 6C. We assume that for the no preview case *H*_*R*_ processes *r*(*t*) and for the preview case a modified reference controller, *H*_*P*_, processes the entire reference preview trajectory *r*_*p*_(*t*) (Fig. 9B). Using this configuration, we can directly model the reference controller as well as the modified reference controller from the output, reference, and disturbance signals using the following equations:

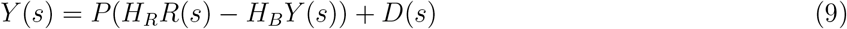

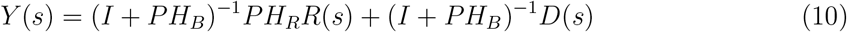

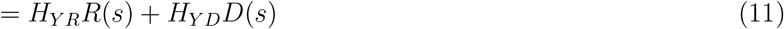

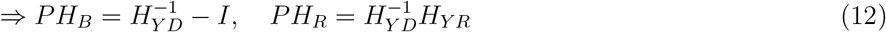

By plugging equation (12) into equations (4) and (5), the new control scheme can be expressed in terms of the previous control scheme using the following equations:

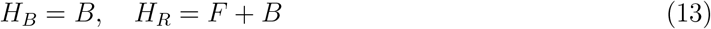

Similarly, for the preview tracking model, we computed *PH*_*BP*_ and *PH*_*P*_ using equation (12) from the output of the preview tracking, reference, and disturbance signals. We assessed the effect of preview on the feedback control pathway by examining the ratio of *PH*_*BP*_ and *PH*_*B*_. The results showed that the ratio was close to 1, indicating that the reference preview does not affect the feedback controller, given the assumption that plant dynamics are unaltered from trial to trial. To investigate the utilization of preview information by subjects, we further evaluated the ratio of *PH*_*P*_ and *PH*_*R*_. The preview reference controller can be expressed as:

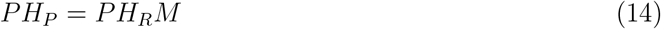

x
where *M* represents ratio of reference controller between preview and no-preview, as illustrated in Fig. 6A. We can view *M* as a reference filter that processes the extra information provided by the preview, i.e. *H*_*P*_ is the no-preview controller (*H*_*R*_) combined with a pre-processing reference filter (*M*). Equation (14) enables us to incorporate the reference filter, *M*, into our original feedforward-feedback control scheme, resulting in the preview control mechanism depicted in Fig. 6C. By comparing the preview filters *M* for ataxia group and control group, we can explore the role of the cerebellum in the utilization of preview information. Additional information on the modeling of the preview filter *M* can be found in the supplementary section 6.2.

### 5.6 Statistical Analysis

All statistical analyses were performed with custom scripts written in MATLAB 2022B (Mathworks, Inc, Natick, MA, USA) using MATLAB’s statistics toolbox. The statistical tests used for each analysis is described in the captions of the associated figures.

### 5.7 Stability Margin Analysis of the Feedback Pathway

Stability margins serve as a measure of the robustness of the feedback control system. The critical juncture where the system approaches instability occurs when the gain is equal to 1 at a phase of 180^°^. The gain margin represents the quantitative measure of the system’s capacity to tolerate variations in gain before it becomes unstable, particularly when the phase is at 180^°^; a higher gain margin implies greater robustness. On the other hand, the phase margin measures the system’s ability to maintain stability in the face of phase shifts, particular near a gain equal to 1. A larger phase margin indicates increased robustness against phase changes.

For formal definitions of stability margins, refer to [48], and for recent applications of stability margins to sensorimotor control, see [49]. In this study, stability margins were calculated for each participant. First, the frequency responses were smoothed by averaging the magnitudes and phases of neighboring frequency points within a specified window size, excluding the first and last frequencies to preserve boundary integrity. The smoothed data was then used to compute the gain and phase margins through linear interpolation.

## 6 Supplementary materials

### 6.1 Model Fitting for Feedforward and Feedback Pathway

We listed the top two model candidates for both the feedforward and feedback pathways in Fig. 10A. We have also plotted the model fitting performance, showing the cross-validation error (CVE) versus the fitting error in Fig. 10B,C.

**Figure 10:**
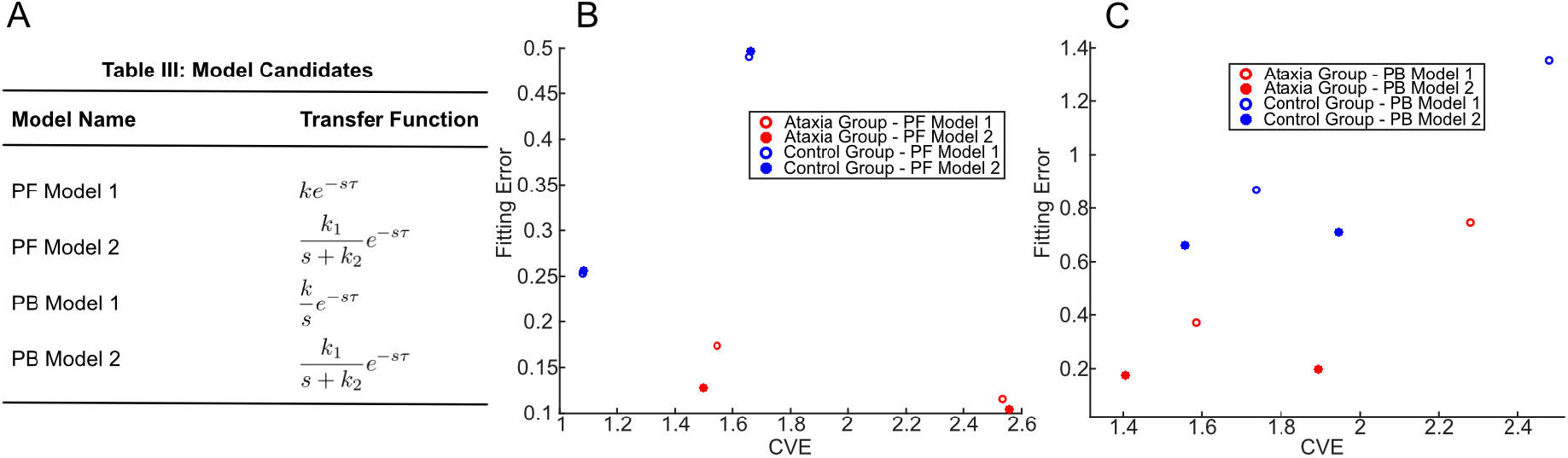
Model Fitting for Feedforward and Feedback Pathway. A. Top two model candidates for the open loop feedforward pathway (*PF*) and open loop feedback pathway (*PB*). B. Model fitting performance for feedforward pathway. The figure displays the fitting error and cross-validation error (CVE), with two data points for each model candidate corresponding to the x-axis and y-axis fitting. PF model 2 closely resembles PF model 1, but with one additional parameter. Therefore, we select PF model 1, a pure-gain-with-time-delay model, i.e. *ke*^−*sτ*^ as the best-fitted model for the feedforward pathway. Same for the control group and ataxia group. **C. Model fitting performance for feedback pathway**. The PB model 2 exhibits smaller fitting error and smaller cross-validation error (CVE). We select PB model 2, a leaky-integrator-with-time-delay model, i.e. 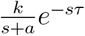, as the best-fitted model for the feedback pathway. Same for the control group and the ataxia group.

### 6.2 Preview Filter Modeling

Using the model fitting techniques described in Methods, we created a best fit model of *M* for both the ataxia group and control group. The same model structure fit both groups, with slightly different parameters (Fig. 11). The model has the following form:

**Figure 11:**
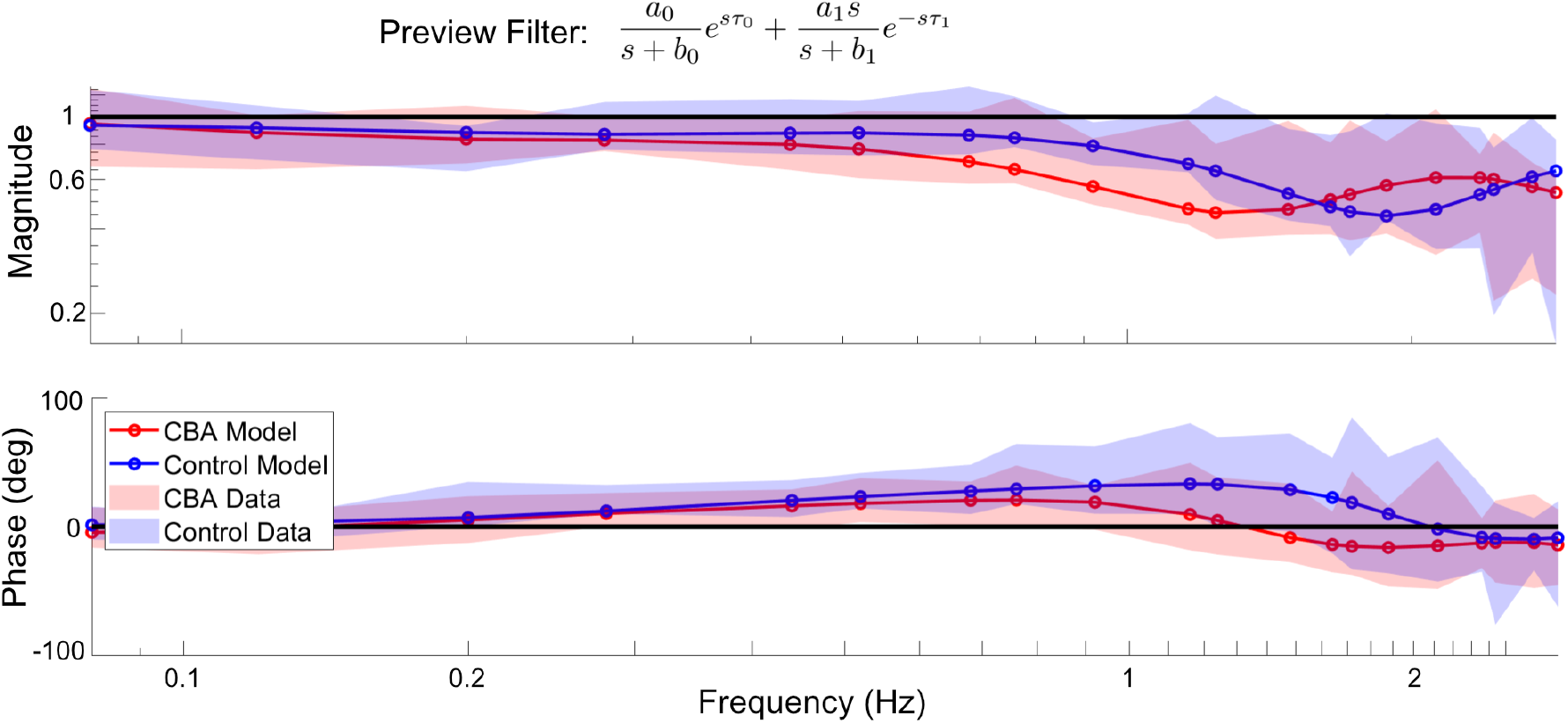
Model of preview filtering. The model of the preview filtering, *M*, introduced in Fig. 6C, fit to the ataxia group’s and control group’s data. The best-fit model is one that combines a low pass filter with a time lead 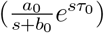 and a high pass filter with a time delay 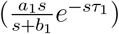. The fitted model for control group is 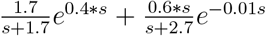, and for ataxia group is 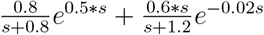.

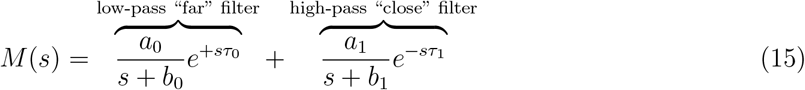

The first term is a low-pass filter (LPF) with time lead *τ*_0_, capturing the fact that participants can use “future” information from the preview provided. The second term is a high-pass filter (HPF) with a time delay *τ*_1_, corresponding to accentuating rapid changes of the actual target.

The structure of the model suggests what pieces of information subjects are using from the reference preview. There are two important components to the model: a time-lead component based on a “future/far point” and a time-lag component based on a “near point”. The time-lead component of the reference filtering is a low pass filter. This suggests that for future points in the preview trajectory, subjects may focus on the low frequency movements, meaning the general motion path of the future points is of interest, not precisely capturing the more rapidly oscillating components of the movement. Conversely, the time-lag component of the reference filtering is a high pass filter. This suggests that for points on the trajectory closer to the reference, there is a focus on precisely capturing the high frequency, seemingly erratic, parts of the movement trajectory. These two components in tandem suggest that subjects are using information from multiple time-points along the reference preview to improve their tracking performance. Note model is phenomenological and highly simplified (only two terms), but is consistent with participants using multiple points along the preview trajectory.

